# Multiple Capsid Protein Binding Sites Mediate Selective Packaging of the Alphavirus Genomic RNA

**DOI:** 10.1101/2020.07.20.212746

**Authors:** Rebecca S. Brown, Dimitrios G. Anastasakis, Markus Hafner, Margaret Kielian

## Abstract

The alphavirus capsid protein (Cp) selectively packages genomic RNA (gRNA) into the viral nucleocapsid to produce infectious virus. Using photoactivatable ribonucleoside crosslinking and an innovative biotinylated Cp retrieval method, we comprehensively defined binding sites for Semliki Forest virus (SFV) Cp on the gRNA. While data in infected cells demonstrated Cp binding to the proposed genome packaging signal (PS), mutagenesis experiments showed that PS was not required for production of infectious SFV or Chikungunya virus. Instead, we identified multiple novel Cp binding sites that were enriched on gRNA-specific regions and promoted infectious SFV production and gRNA packaging. Comparisons of binding sites in cytoplasmic vs. viral nucleocapsids demonstrated that budding caused discrete changes in Cp-gRNA interactions. Notably, Cp’s top binding site was maintained throughout virus assembly, and specifically bound and assembled with Cp into core-like particles *in vitro*. Together our data suggest a new model for selective alphavirus genome recognition and assembly.

## INTRODUCTION

Alphaviruses are enveloped, highly organized RNA viruses that include a number of important human pathogens such as Chikungunya virus (CHIKV), Ross River virus (RRV), and Venezuelan Equine Encephalitis virus (VEEV) ^1^. These viruses are transmitted by mosquito vectors and can infect a wide variety of mammalian and avian species. Alphaviruses are organized into complexes based on genetic and antigenic relationships ^2^. While the general features of structure and lifecycle are shared, alphaviruses can differ in properties such as receptor usage, tissue tropism and pathogenesis. Within the Semliki Forest virus (SFV) complex, CHIKV and RRV cause severe polyarthritis and have emerged to produce epidemics that affect millions of people globally ^3–5^. To date, there are no specific antiviral treatments or licensed vaccines for any alphavirus.

Alphaviruses infect host cells by endocytic uptake and membrane fusion, thereby delivering the nucleocapsid (NC) core into the cytoplasm where replication takes place^1^. NC disassembly is thought to be mediated by interactions with ribosomes, resulting in the uncoating of the RNA genome ^6^. The viral genome (gRNA) is an ∼11.5 kb single-stranded plus-sense RNA that is 5’-capped and contains a 3’-polyA tail. It is translated to produce the viral nonstructural proteins (nsP1-4), which assemble a membrane-associated replication complex (reviewed in ^1,7^). An antisense complement of the gRNA is synthesized and this negative-sense RNA serves as the template for producing new gRNAs. In addition, an internal promoter drives the production of a smaller RNA that is identical in sequence to the last ∼1/3 of the gRNA. This subgenomic RNA (sgRNA) is translated to produce the viral structural proteins: capsid protein (Cp), 6K/TF, and the E1 and E2 envelope proteins. Cp specifically interacts with the gRNA to assemble the NC (Fig. 5a). Budding of progeny virus at the plasma membrane releases particles with icosahedral symmetry, containing 240 copies each of Cp, E1, and E2, and 1 gRNA ^1,8^.

Alphaviruses have a robust growth cycle and produce particles with high specific infectivity, indicating strong quality control of particle assembly and genome packaging. However, it is not clear how the specific recognition and assembly of the viral RNA into NC occurs, given that Cp must select gRNA in the presence of a 3-4-fold molar excess of sgRNA produced from the replication complexes ^9–12^. In addition, while virus infection inhibits transcription of most cellular RNAs ^13,14^, abundant host RNAs still remain in the infected cell. Serial passage under high multiplicity conditions can lead to the production of defective viral genomes that can be packaged into defective interfering (DI) particles. Such DI particles are missing portions of the gRNA, and are dependent on wild type helper virus for their propagation. Mapping, sequence analysis and mutagenesis studies of DI particles support the presence of a packaging signal (PS) in the gRNA that promotes its recruitment into virus particles (reviewed in ^15^). The nature and location of the PS appear to vary depending on the virus complex, with the VEEV complex and Sindbis virus (SINV) complex having a PS located in the nsP1 region ^16,17^. In contrast, the PS for the SFV complex including CHIKV and RRV maps within the nsP2 region ^18,19^. The evidence for a role of the PS in genome packaging is strongest for VEEV, where mutation of the PS in nsP1 leads to an approximately 2-log reduction in virus titer while still supporting growth to ∼10^8^ PFU/ml ^17^.

While the inference is that the PS functions by interaction with the Cp, as yet there is no evidence for Cp’s direct binding to the PS in infected cells. Such studies are complicated by the fact that the highly basic nature of the Cp promotes its non-specific interaction with nucleic acids and proteins *in vitro* or in cell lysates ^6,20–24^. Here we developed a novel biotinylation system to stringently and efficiently retrieve the capsid protein from SFV-infected cells or virus particles. We combined this with PAR-CLIP (photoactivatable ribonucleoside crosslinking and immunoprecipitation) ^25^ to comprehensively map Cp interactions on the SFV genome with nucleotide precision. Our results identify novel Cp binding sites on the gRNA, rule out a role for the SFV and CHIKV PS, highlight changes in Cp-gRNA interactions during virus biogenesis, and identify a site on the gRNA that Cp preferentially binds and assembles with *in vitro*. Together our data support a new model for alphavirus gRNA packaging and assembly that is mediated by Cp binding to multiple sites on the gRNA.

## RESULTS

### AVI-tagged Cp is a new tool to stringently and efficiently retrieve Cp

The alphavirus Cp is composed of an N-terminal polybasic RNA-binding domain that mediates gRNA packaging, and a C-terminal protease domain that forms the outer NC shell (reviewed in ^15^). The charged nature of the Cp (pI ∼ 10) promotes non-specific, electrostatically-driven interactions, but wash conditions sufficient to disrupt them lead to poor protein retrieval by a variety of polyclonal and monoclonal Cp antibodies (RSB, unpublished observations). Moreover, prior studies showed that tagging Cp with proteins such as mCherry caused aberrant Cp localization and NC morphology (e.g., ^26^). In order to map Cp’s binding sites on the gRNA, we devised a novel Cp retrieval strategy that takes advantage of the high affinity interaction of biotin with Streptavidin.

The 13 residue minimal biotin acceptor peptide (mAVI tag; Fig. 1a) is specifically and efficiently biotinylated by the biotin ligase BirA ^27–30^. We engineered the mAVI tag into the SFV infectious clone after Cp residue 95 (SFV-mAVI; Fig. 1a), a site previously shown to be permissive for small peptide insertions ^26,31^. As a host cell for our studies we constructed a Vero cell line stably expressing humanized BirA in the cytoplasm (Vero BirA cells). Virus growth experiments showed that the mAVI tag was well tolerated, causing a modest decrease in maximal titers (Fig. S1a). We observed no difference in virus or NC morphology by transmission electron microscopy (Fig. S1b), or in the expression or localization of biotinylated Cp-mAVI vs. Cp-WT in infected cells (Figs. 1b, S1c-e.) No change was detected in SFV-mAVI vs. WT specific infectivity as calculated by comparing the number of infectious particles to total particles (PFU:E2 ratio) (Fig. 1c), thus indicating accurate packaging of the gRNA by Cp-mAVI. Using the biotin handle on Cp, we optimized retrieval conditions with Streptavidin beads to efficiently and specifically pull-down Cp. Under these stringent conditions (including 0.5% SDS and washing with 0.5 M NaCl), Cp was retrieved from infected BirA cells only when biotin was present in the culture medium (Fig. 1d).

**Figure 1.**
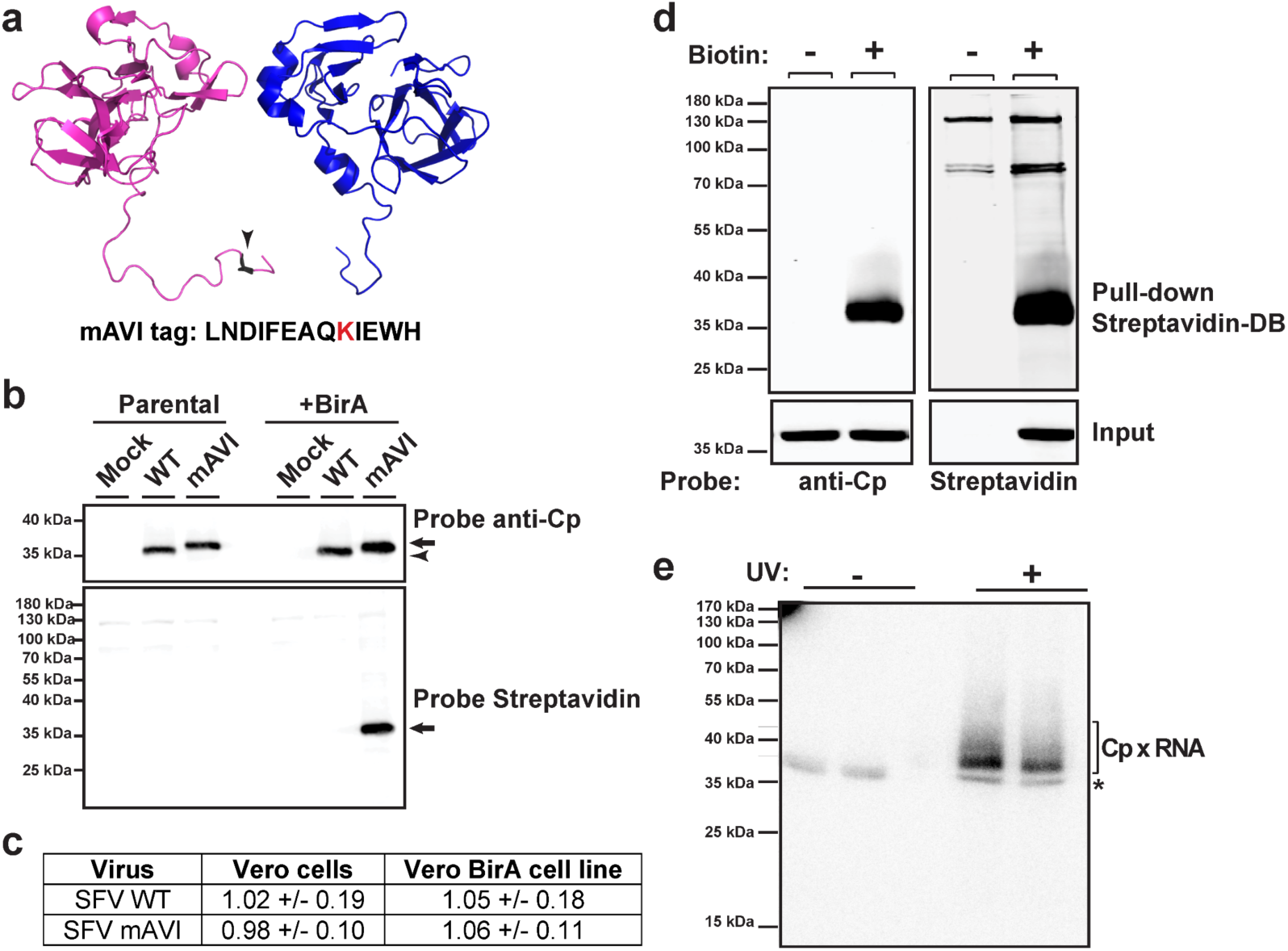
Properties and retrieval of SFV Cp using mAVI tag. (a) Partial structure of Eastern Equine encephalitis virus Cp dimer encompassing (on the left) Cp residues 85- 261 (PDB: 6MX7) ^59^. Arrowhead indicates the homologous location where the mAVI tag was inserted in SFV Cp. The highlighted lysine residue within the mAVI tag sequence is the target for BirA biotinylation. (b) Vero parental or Vero BirA (+BirA) cells were mock- infected, infected with SFV WT, or infected with SFV mAVI. At 7.5 hpi, lysates were harvested, subjected to SDS-PAGE, and analyzed by western blot using a polyclonal antibody against Cp or a Streptavidin Alexa-680 probe. Arrowhead indicates Cp WT, and arrows indicate Cp-mAVI. (c) Specific infectivity was measured as the ratio of the number of infectious particles (determined by plaque assay) to the number of total particles (determined by quantitative western blot of purified virus using an E2-specific antibody). Data are the average and range from two independent experiments. Units represent ratio of PFU/E2 signal. (d) Vero BirA cells were grown in biotin-depleted media for 3 days (-biotin) or grown in normal media (+biotin). Cells were infected with SFV mAVI, and lysates were harvested at 7.5 hpi, retrieved with Streptavidin DynaBeads (SA-DB), and analyzed by SDS-PAGE and western blot using a monoclonal antibody against Cp or a Streptavidin Alexa-680 probe. (e) Autoradiogram of Cp-RNA crosslinked adducts retrieved with SA-DB from Vero BirA cells infected with SFV mAVI (representative of n=2 independent experiments). Asterisk indicates an infection- and UV-independent band (Figure S1e). See also Figure S1.

### Cp binds many discrete sites across the genome

To define the Cp binding sites on the gRNA in infected cells at nucleotide resolution, we combined our optimized Cp retrieval system with the sensitive and specific PAR-CLIP method ^25^. PAR-CLIP uses a photoactivatable ribonucleoside to crosslink labeled RNAs to their binding proteins, with up to 1000-fold higher crosslinking efficiency compared to traditional CLIP-seq approaches. Crosslinking induces T-to-C mutations during cDNA library generation, enabling precise identification of a protein’s specific binding sites on the RNA ^25^. Vero BirA cells were infected with SFV-mAVI for 1.5 h. The cells were then cultured in the presence of excess biotin to ensure robust Cp-mAVI biotinylation and the photoactivatable ribonucleoside 4-thiouridine (4SU) to label nascent RNAs. 4SU had no effect on virus growth (data not shown). Because alphaviruses inhibit Pol II transcription early in infection ^13,14^, this 4SU labeling strategy strongly biases labeling towards viral RNAs over cellular RNAs. At 7 hours post-infection cells were irradiated with UV (λ >310 nm) to crosslink RNAs with bound proteins, lysed, and RNAs digested with RNase T1 to produce footprints protected by RNA binding proteins. The total cellular pool of Cp-mAVI-biotin was then retrieved with Streptavidin beads, and crosslinked RNAs were 5’-end labeled with γ-^32^P-ATP and subjected to SDS-PAGE followed by transfer to a nitrocellulose membrane. The resulting Cp-RNA adducts were only detected upon UV irradiation and were the only UV-dependent crosslinked products that were retrieved (Figs. 1e and S1f).

The RNAs crosslinked to Cp were purified and converted into cDNA libraries and sequenced using the Illumina MiSeq Platform (see Methods for details). From two biological replicates we obtained 1,384,633 and 3,213,621 sequence reads of which 121,119 and 284,837 mapped to the viral genome, respectively. For further analysis we only considered the 105,920 and 233,188 sequence reads, respectively, that contained the diagnostic T-to-C mutation introduced during cDNA library construction of 4SU- labeled and crosslinked RNA. This allowed us to a) remove background sequences from co-purifying, non-crosslinked fragments from abundant RNAs and b) identify the crosslinking site at nucleotide resolution. Comparison of the crosslinked sequence reads revealed an excellent correlation for read density of the gRNA between the two biological replicates (Pearson correlation coefficient r=0.8123). We proceeded to combine the two datasets and count normalized read density across every nucleotide in the gRNA (Fig. 2a). Cp binding sites were defined using an arbitrary cutoff of read density (Experimental Methods). Altogether we defined 58 high-confidence sites with a median length of 24 nt (Table S1). Approximately ∼100 additional Cp interaction sites could be observed on the gRNA, but with significantly lower read densities (Table S2).

**Figure 2.**
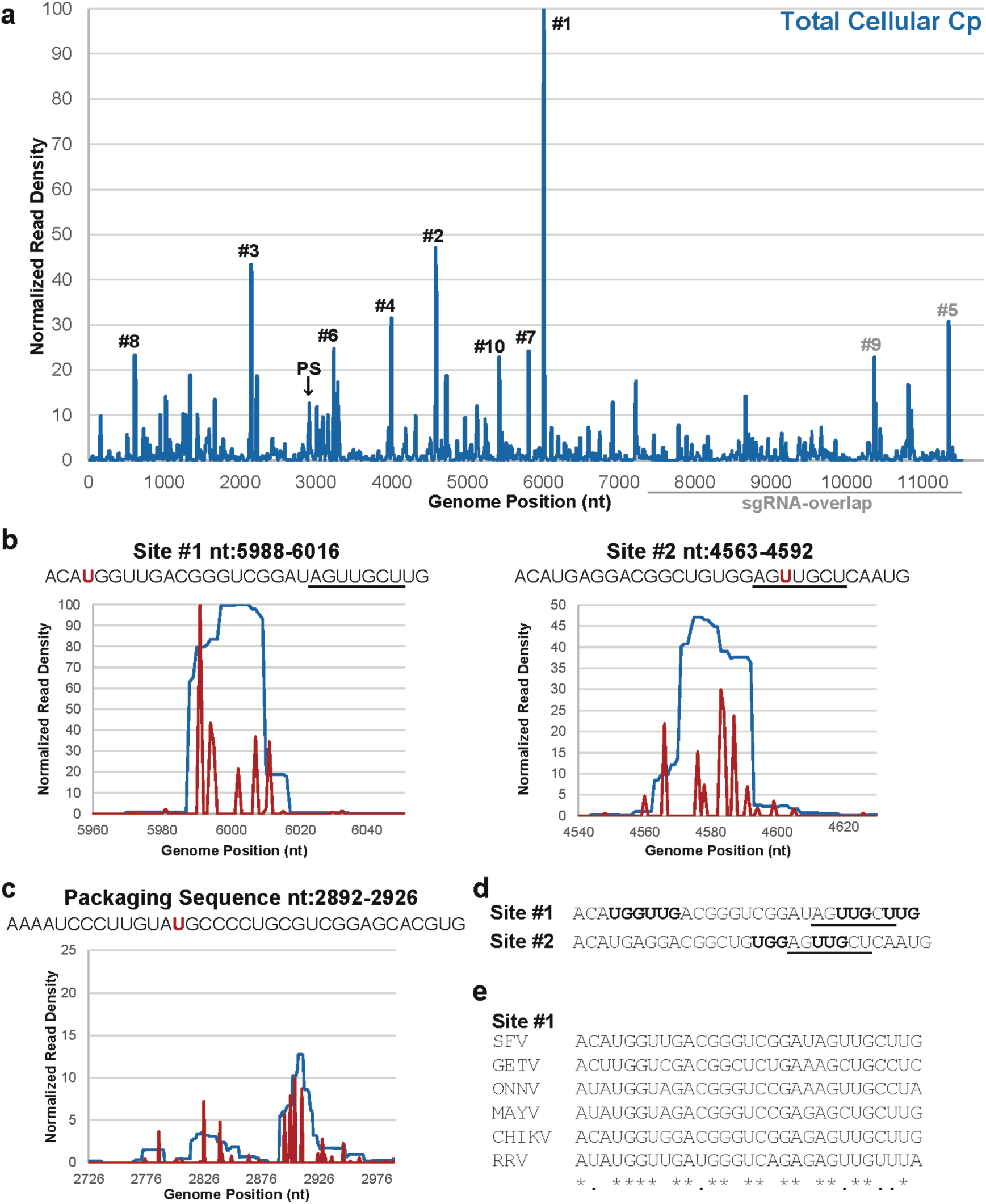
Cp’s genomic RNA binding sites in cells. (a) Normalized read density from two independent PAR-CLIP experiments plotted across genome nucleotide position. Reads were normalized to the nucleotide position with the highest read frequency within the library, which were read values of 14,134 and 13,343, before averaging. Arrow indicates Cp’s specific binding site within the Full PS, #1 indicates site #1, #2 indicates site #2, etc. Gray line represents the sgRNA-overlapping region of the gRNA. (b) Normalized read density (blue lines) across site #1 (nt:5988-6016; left) and site #2 (nt4563-4592; right), and the normalized read frequencies of the T-to-C mutations (red lines). T-to-C frequencies were normalized to the T with the highest C mutation frequency within each library replicate, which were frequencies of 5,386 and 4,750, before averaging. The T/U residue with the most C mutations for each site is highlighted in red within their respective sequence. The 7 identical nucleotides between site #1 and 2 are underlined. (c) As in (b), but for the Full PS (nt:2726-2991). The sequence corresponding to the Cp-PS binding site (nt:2892-2926) is shown at the top, with the T/U residue with the most C mutations highlighted in red. (d) Site #1 and #2 sequences. Bolded nucleotides are either “UUG” or “UGG” motifs. The 7 identical nucleotides between site #1 and 2 are underlined. (e) Sequence alignment of site #1 with the homologous sites in SFV complex members GETV (AY702913.1), CHIKV (EF452493.1), MAYV (AF237947.1), ONNV (M20303.1), and RRV (GQ433359.1). Asterisks indicate nucleotide identity, and periods indicate nucleotide similarity. See also Figure S2, Figure S3, Figure S4, Table S1, and Table S2.

Analysis of the nucleotide composition of Cp’s top 58 binding sites revealed that these sequences were relatively U-rich and A-poor (Fig. S2a). In contrast, sequences negative for Cp binding in either replicate were relatively A-rich (Fig. S2a). Previous studies show that PAR-CLIP can identify binding sites of all compositions. For example, binding sites for the HIV-Gag protein are not U-rich ^32^, while CNBP and DHX36 selectively bind G-rich sites ^33,34^. Thus our results suggest that the SFV Cp does in fact preferentially bind U-rich sequences in the cell. A computational search ^35^ for sequence motifs enriched in the top 58 Cp binding sites revealed that GC- and UG-based motifs were common, but did not identify a specific sequence motif from these 58 sites (Fig. S3a).

The published DI-RNA studies mapped the SFV PS to a 266 nt region located within the nsP2 gene (nt:2726-2991; here termed Full PS), suggesting a role for this region in gRNA packaging ^18,19^. Our PAR-CLIP data showed that Cp bound a 35 nt region towards the 3’ end of this region (nt:2892-2926; here termed Cp-PS binding site), with weaker Cp binding also observed within nt:2815-2856 (Fig. 2c). This is the first direct demonstration of in cell interaction of Cp within the Full PS region. However, the Cp-PS binding site was not the top binding site on the gRNA, with 20 other sites showing higher binding (Fig. 2a and 2b).

We re-analyzed the Cp binding sites on the gRNA using the Cp-PS binding site as a lower cutoff. Within these 21 high affinity sites there was a stronger trend towards Cp binding U-rich and A-poor sequences (Fig. S2a). Sequence motif analysis of the top 21 sites showed an enrichment for UUG and UGG-trinucleotide motifs (Fig. S3a). Secondary structure predictions for the top 21 sites predicted all 21 RNAs would adopt stem-loop structures, but of varying sizes (Fig. S3b)^36^.

Manual inspection of Cp’s highest affinity sites revealed that Cp’s top two binding sites (nt:5988-6016 and nt:4563-4592) (Fig. 2b, sites #1 and #2) are very similar in sequence and contain UUG and UGG motifs (Fig. 2d, bolded). Furthermore, sites #1 and #2 share an identical stretch of 7-nucleotides not present at any other location within the gRNA (Fig. 2b and d, underlined). Sequence alignment analysis showed that site #1’s sequence is hyperconserved across several other viruses in the SFV complex compared to their overall sequence identity (Fig. 2e), while site #2 and twelve other top binding sites are hyperconserved in CHIKV as well (Fig. S4a). Closer inspection of site #1 and #2’s predicted secondary structures revealed similar stem-loop structures containing tandem G:U wobble base pairs within the stem (Fig. S3b). Tandem G:U wobble base pairs were also predicted to form in the stems of sites #8, #14, and #15 (Fig. S3b).

### Cp preferentially binds sequences unique to the genomic RNA

The sgRNA is identical in sequence to the last ∼1/3^rd^ of the gRNA and is present in 3-4X molar excess over it in infected cells ^9–11^. It has therefore been hypothesized that Cp binding sites specific to the gRNA, such as the PS, dictate selective packaging in infected cells. Inspection of our PAR-CLIP data suggested that Cp binding was biased towards the first ∼2/3^rd^ of the genome (Fig. 2a). To formally test this, we summed the number of observed reads at each nucleotide position and categorized them as mapping to the gRNA-specific region vs. mapping to the sgRNA-overlapping region (Fig. 3a). We performed a chi-square analysis to test whether the observed summed read distributions were statistically different from our null hypothesis, in which random Cp binding would result in ∼2/3 of the summed reads mapping within the gRNA-specific region and ∼1/3 within the sgRNA region. The results showed that for each replicate, observed read distributions mapping to gRNA-specific regions were significantly higher than expected (p<0.0001) (Fig. 3a). This result was not due to a nucleotide bias as the gRNA-specific and sgRNA sequences have nearly identical compositions (Fig. 3b). The enriched binding of Cp to gRNA-unique sequences suggests a direct molecular rationale for selective Cp recognition and packaging of gRNA vs. sgRNA.

**Figure 3.**
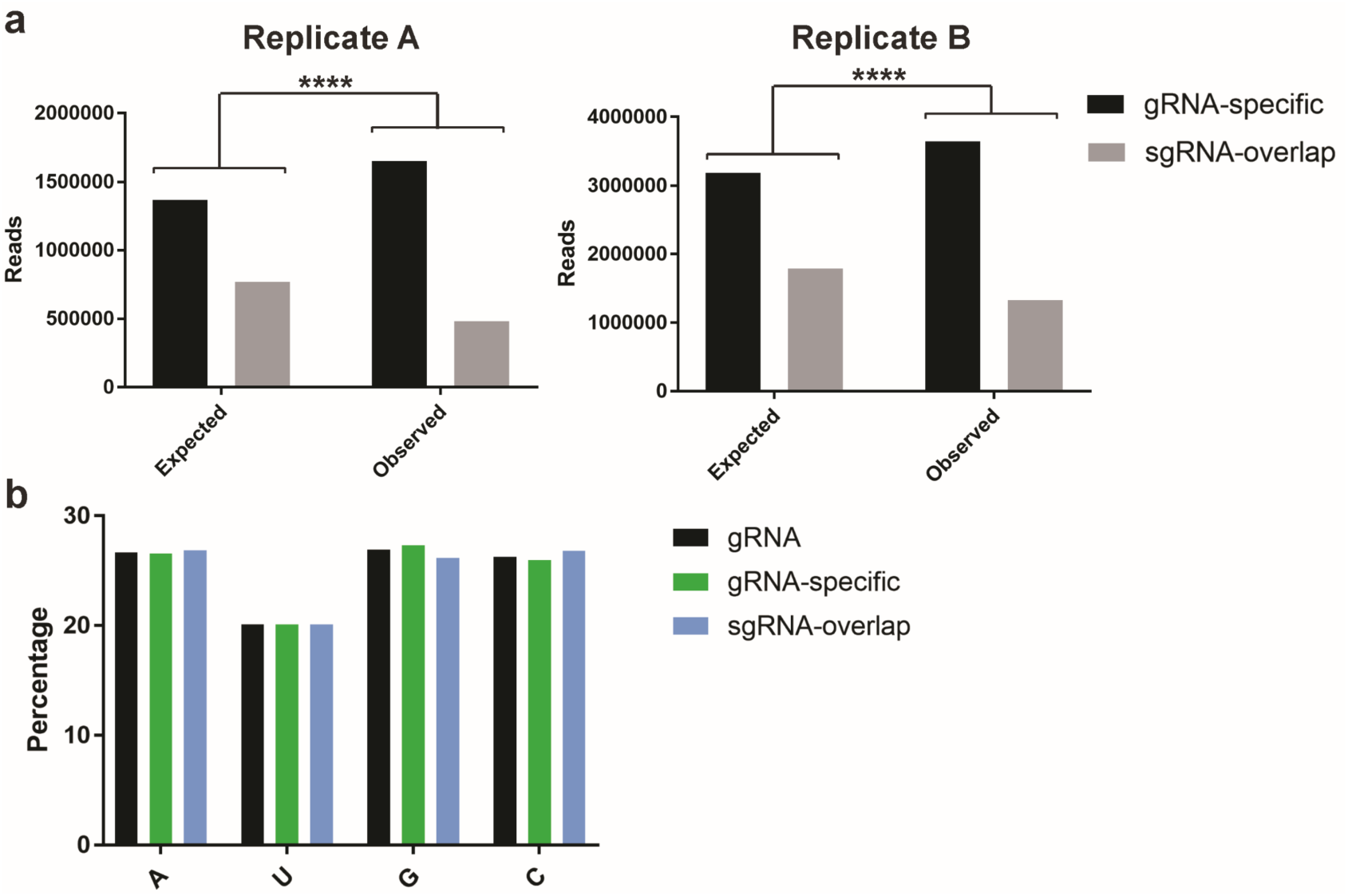
Cp preferentially binds genome-specific sequences. (a) Chi-square analysis test for replicate libraries (Rep A and Rep B) comparing the expected vs. observed number of sequence reads summed per nucleotide position mapping to gRNA-specific sequences (black) or sequences that overlap with the sgRNA (gray). Two-sided, df=1, **** indicates p<0.0001. (b) Nucleotide composition of the gRNA, gRNA-specific sequences, and sequences that overlap with the sgRNA.

### Mutation of multiple Cp binding sites inhibits infectious virus production and rules out a role for the PS

We set out to assess the significance of Cp’s top binding sites on genome packaging by mutating them and measuring the effect on infectious virus production. All mutations were designed to strictly maintain amino acid sequence identity, disrupt predicted secondary structures, and minimize rare codon usage and changes in dinucleotide frequencies (Table S3). We first mutated the Full PS previously identified by DI RNA studies (see Fig. S3c for predicted effects on PS secondary structure). To our knowledge, while such mutational analyses have been performed for the SINV and VEEV PS ^17^, this is the first direct test of PS function for any SFV complex virus. We infected Vero cells with the SFV WT or Full PS mutant and measured infectious virus production over time. The Full PS mutant did not affect virus growth compared to WT (Fig. 4a). This is consistent with our PAR-CLIP data, which demonstrated that Cp does not extensively interact with the PS region, and more efficiently binds ∼20 other sites on the gRNA.

**Figure 4.**
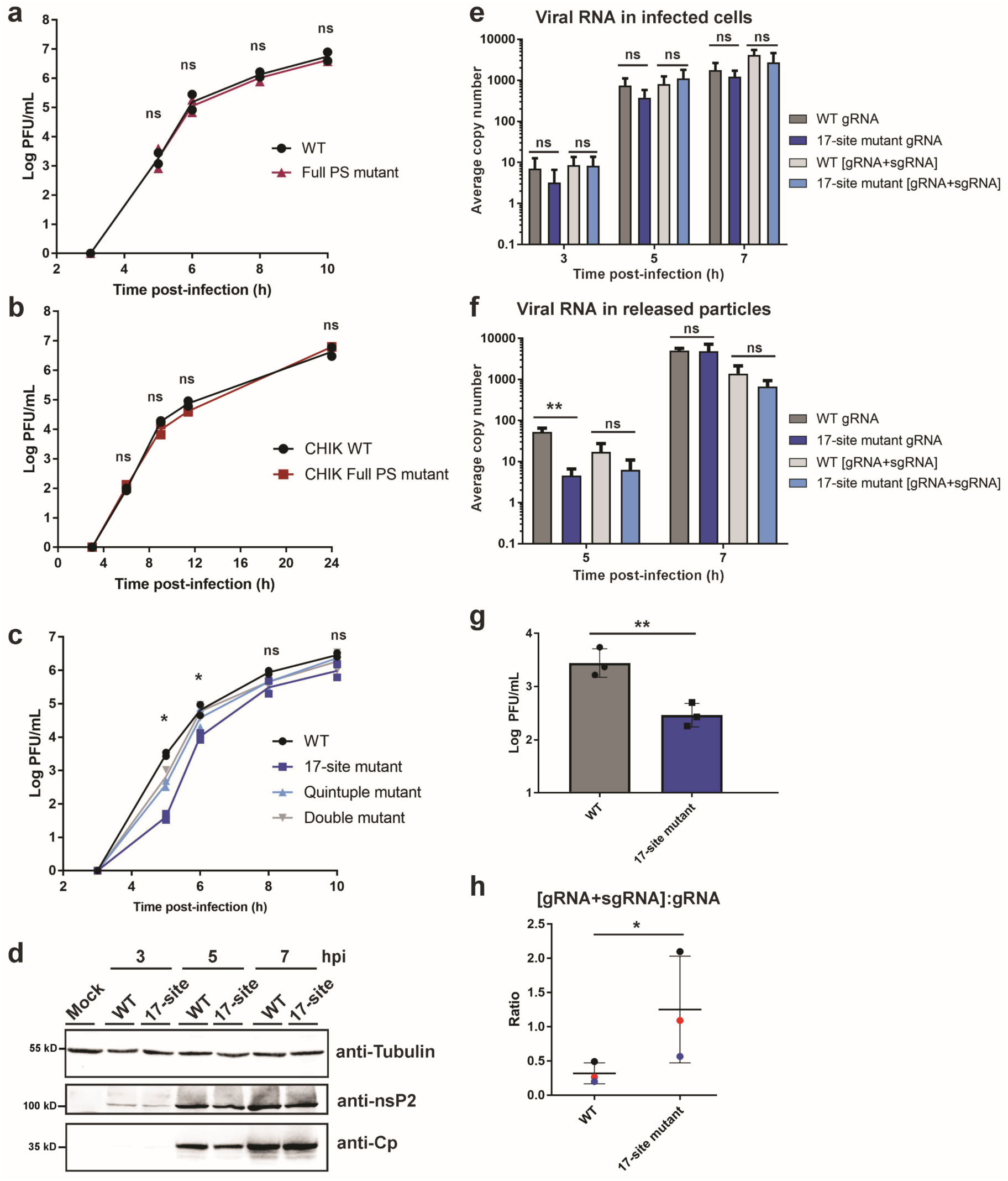
Mutation of multiple Cp binding sites inhibits infectious virus production independent of the PS. (a) Vero cells were infected with SFV WT or Full PS mutant at MOI=0.01. Media from the indicated time points were titered by plaque assay. Individual points from n=2 were plotted. Student’s t-tests compared WT vs. Full PS mutant; ns denotes no significant difference. (b) As in (a), but with CHIKV WT and CHIKV Full PS mutant at MOI=0.01. (c) As in (a) and (b), but with SFV WT, 17-site mutant, Quintuple mutant (sites #1-4 and Cp-PS binding site), and Double mutant (sites #1 + 2). Student’s t-tests compared WT vs. 17-site mutant; * denotes p<0.05. (d) Lysates from Vero cells mock infected, or infected with SFV WT or the 17-site mutant at MOI=10 were analyzed by western blot for tubulin, nsP2 or Cp. Representative images of n=2. (e) Total cellular RNA was extracted from Vero cells infected with SFV WT or 17-site mutant at MOI=0.01. gRNA and [gRNA+sgRNA] copy numbers in infected cells were determined by RT-qPCR and represent the average per 5 ng of total RNA (n=3). Student’s t-tests compared copy numbers. (f) Media from Vero cells infected with SFV WT or 17-site mutant at MOI=0.01 were pelleted to isolate virus particles. gRNA and [gRNA+sgRNA] copy numbers were determined as in (e) and represent the average copy number per 10% of the pelleted samples (n=3). Student’s t-tests compared copy numbers; ** denotes p<0.01. (g) SFV WT and 17-site mutant samples from the 5 h time point in (f) were titered by plaque assay. Error bars represent standard deviation. Student’s t-test compared WT and 17-site mutant titers; ** denotes p<0.01. (h) The ratio of [sgRNA+gRNA] to gRNA levels from the 5 h time point in (f) for SFV WT and 17-site mutant. The thick black horizontal bar represents the mean and the thin horizontal bars represent the standard deviation. Paired replicates are plotted in the same color. Student’s t-test compared the ratios; * denotes p<0.05. See also Figure S4 and Table S3.

To test if other alphaviruses in the SFV complex were dependent on the previously identified PS, we mutated the region of the CHIKV gRNA corresponding to the SFV DI-RNA-derived Full PS (Fig. S4b and Table S3). Similar to our results with SFV, we observed no effect of mutation of the CHIKV Full PS on infectious CHIKV production (Fig. 4b).

We next focused on a series of mutational analyses of the top gRNA-specific binding sites. Mutation of Cp binding sites #1 and #2 (Double mutant) had no significant effect on infectious virus production (Fig. 4c). Mutagenesis of sites #1-4 plus the Cp-PS binding site (Quintuple mutant) caused a small but statistically significant growth defect at 5 hpi (Fig. 4c); growth at later time points was comparable to that of WT. We then tested if packaging could occur through cooperation among multiple Cp binding sites on the gRNA. Such a multi-site packaging mechanism has been described for some plus-sense RNA viruses ^15,37^. Using the 35 nt Cp-PS binding site as a threshold, we mutated 17 gRNA-specific binding sites including Cp-PS, representing Cp’s highest affinity binding sites on the gRNA (Table S3). Other than the site within the PS, none of these have been previously implicated in any stage of the alphavirus life cycle. The 17-site mutant produced significantly less infectious virus compared to WT (p<0.05) at early time points of virus production (Fig. 4c; 5 and 6 hpi). Combining the 17-site mutant with the full PS mutant produced no additional decrease in virus production (data not shown). The 17-site mutant phenotype became more moderate at late times of virus production (Fig. 4c; 8 and 10 hpi), suggesting that these sites are most important at early time points in virus assembly when Cp levels are limiting. This is consistent with a multi-site packaging mechanism where high affinity binding sites on the gRNA have a greater impact on gRNA recognition and NC assembly at early infection times when Cp levels are lower ^38^.

To determine if the 17-site mutant specifically impacted gRNA packaging and virus assembly vs. aspects of virus replication, we first checked viral protein expression levels by western blot. We observed comparable levels of both nsP2 and Cp between WT vs. 17-site mutant-infected cells (Fig. 4d). We next used RT-qPCR to measure cellular gRNA and sgRNA levels over the course of virus infection, using primers that either specifically amplify the gRNA or amplify both gRNA and sgRNA [gRNA+sgRNA] due to their overlapping sequences. No significant difference was observed in cellular gRNA or [gRNA+sgRNA] levels between WT and 17-site mutant-infected cells from 3-7 hpi (Fig. 4e). Thus, although the gRNA mutations significantly reduce infectious virus production, this was not due to defects in viral RNA replication or protein expression.

We next asked whether the reduction in infectious particles could be due to a decrease in genome packaging in the 17-site mutant. We infected cells, purified SFV WT or 17-site mutant viruses, and quantitated viral gRNA and [gRNA+sgRNA] levels in the released particles. There was a significant decrease in gRNA in the 17-site mutant virus collected at 5 hpi (>10-fold reduction; p<0.01) (Fig. 4f). There was no significant difference in gRNA levels at 7 hpi, or in [gRNA+sgRNA] levels for either time point (Fig. 4f). The ∼10-fold reduction in gRNA levels in the 17-site mutant virus at 5 hpi correlated well with the ∼10-fold reduction in infectious particles observed in a parallel sample at 5 hpi (Fig. 4g).

At these time points and multiplicities we were unable to directly quantitate particle numbers by quantitating E2 in released viral particles (data not shown). Since it was previously demonstrated that a decrease in alphavirus gRNA packaging can result in increased packaging of the sgRNA ^11,17^, we analyzed if there was more sgRNA packaged into 17-site mutant particles. Results showed that even though the gRNA levels were significantly reduced in the mutant virus particles at 5 hpi, their [gRNA+sgRNA] levels were not significantly different from those of WT virus (Fig. 4f). Comparing the ratio of [gRNA+sgRNA] to gRNA between the WT and 17-site mutant particles showed a significantly higher ratio (∼4-fold, p<0.05) for the 17-site mutant compared to WT (Fig. 4h), suggesting a higher proportion of sgRNA in the mutant particles. These data thus suggest that the decrease in gRNA and infectious particles observed for the 17-site mutant is at least in part caused by increased non-specific packaging of the sgRNA.

Together, our results rule out a significant role of the previously proposed DI- RNA-derived PS in infectious virus production for the SFV complex. Instead, they support a mechanism in which alphaviruses package their gRNAs through Cp interaction with multiple high affinity sites present in the gRNA-specific region of the genome.

### Changes in Cp:gRNA interactions during virus exit

Our PAR-CLIP results were obtained by analysis of the total cellular Cp in infected cells, a pool that includes both unassembled Cp and assembled NC (Fig. 5a). To examine different stages of virus assembly, we compared these total cellular Cp results with PAR-CLIP analyses performed on cellular and viral NC produced in 4SU-labeled cells (Fig. 5a). For cellular NC, UV-crosslinking was performed on infected cells as above, followed by cell lysis and subsequent NC purification by sucrose gradient sedimentation (Fig. S2b). For viral NC, released virus particles were pelleted, UV-crosslinked, and then lysed. Biotinylated Cp-RNA adducts from viral and cellular NCs were retrieved with Streptavidin beads (Fig. S2c), as described above. As observed for the total cellular Cp, retrieval was strictly dependent on UV irradiation (Fig. 5b). PAR-CLIP libraries were generated from two biological replicates for each sample type, and reads were processed as above. Biological replicates correlated strongly (Pearson correlation coefficients of r=0.8655 for cellular NC and r=0.9459 for viral NC) and thus we averaged the duplicate libraries and plotted normalized read density across the gRNA sequence (Fig. 5c and 5d).

**Figure 5.**
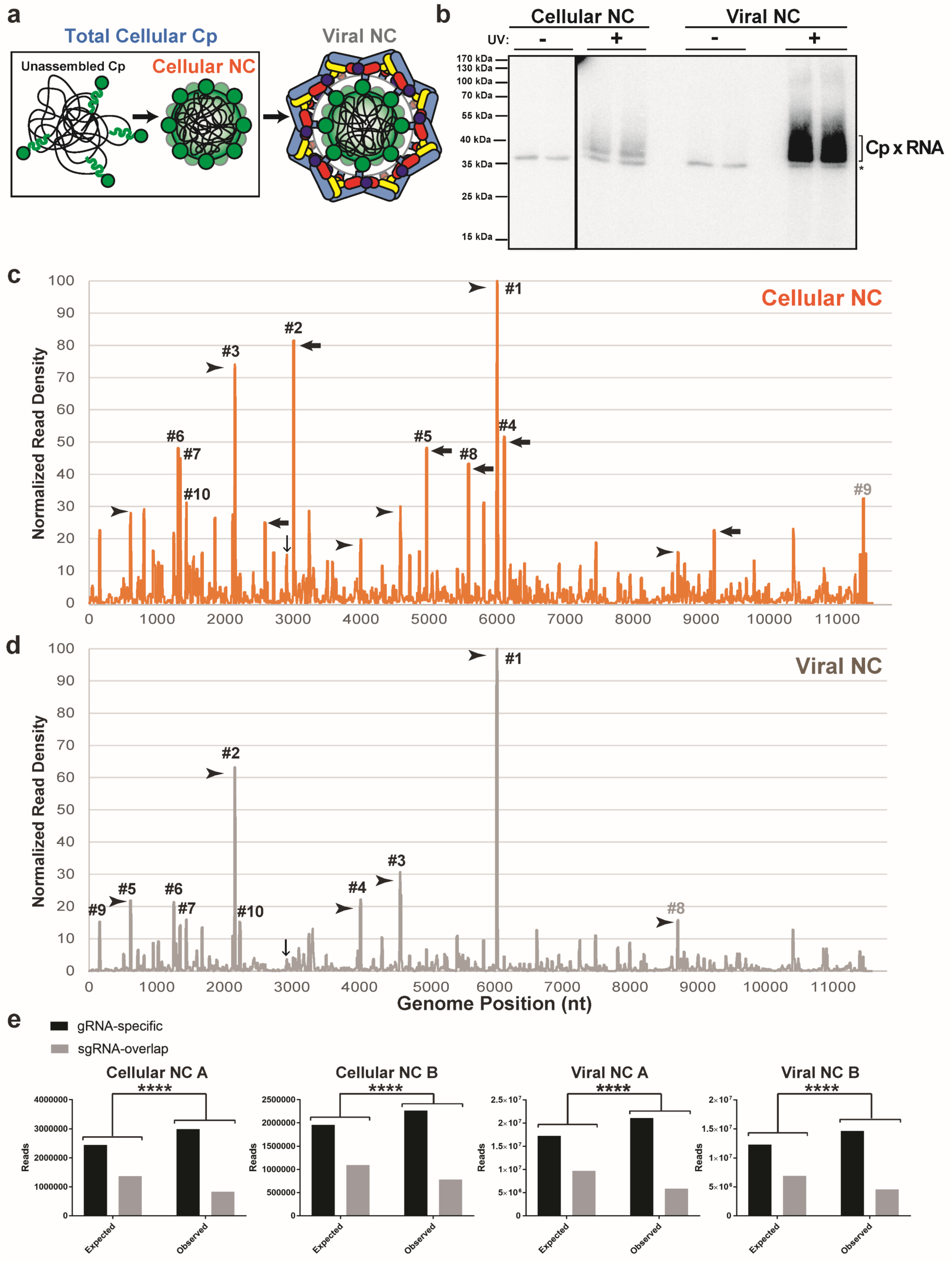
Changes in Cp-gRNA interactions triggered by virus budding. (a) Schematic of the different Cp and NC populations present during virus assembly. Cp (green), gRNA (black), E2 (pale blue), and E1 (colored by domain: dIII blue, dII yellow, and dI red). (b) Autoradiogram of Cp-RNA crosslinked adducts as in Figure 1e, but for purified cellular NC and viral NC. (c) and (d) Normalized read densities as in Fig. 2a, but for cellular NC (c) and viral NC (d). Reads were normalized to the nucleotide position with the highest read frequency within each library, which were read values of 10,143 and 5,855 for cellular NC and 140,546 and 88,226 for viral NC, before respectively averaging. Downward-facing arrow indicates Cp’s specific binding site on the PS, and numbered sites represent the top 10 sites bound for each respective sample type. Arrowheads indicate examples of Cp binding sites maintained throughout virus assembly, and leftward-pointed arrows indicate examples of distinct Cp interactions enriched only in the cellular NC assembly state. (e) Chi-square analysis test as in Figure 3a, but for cellular NC and viral NC replicate libraries. Two-sided, df=1, **** indicates p<0.0001. See also Figure S2 and Table S2.

Overall, Cp:gRNA interactions appear similar across virus assembly states, with many of the same binding sites maintained over the course of assembly (Fig. 5c and 5d, arrowheads). Cp’s highest affinity site in the total cellular Cp sample (Fig. 2b, #1) is also the highest affinity site in cellular NCs (Fig. 5c, termed NC#1) and viral NCs (Fig. 5d, termed V#1), demonstrating that Cp preferentially binds this specific RNA sequence across virus assembly states. Additionally, the third best bound site in the total cellular Cp population (Fig. 2b, #3) was also the third best bound site in cellular NC (Fig. 5c, NC#3) and the second-best bound site in viral NC (Fig. 5b, V#2). Cp’s top four binding sites in the total cellular Cp population also showed strong binding within cellular NC and viral NC (Figs. 2b, 5c, 5d and Table S2), further emphasizing Cp’s affinity for these RNA sequences. In addition, all of the assembly states displayed enriched binding of Cp to gRNA-specific sequences (Fig. 5d).

Distinct changes were observed between cellular NC and viral NC (Fig. 5c, black arrows). For example, Cp’s second highest affinity site in cellular NC is located at nt:2997-3016 (Fig. 5c, NC#2). Upon virion budding, this binding site drops below the top 50 binding sites within the viral NC. This pattern of strong Cp:gRNA binding events in cellular NC that become less prominent in viral NC is also observed for cellular NC sites NC#4 (nt:6096-6115), NC#5 (nt:4954-4972), and NC#8 (nt:5569-5706), among others (Fig. 5c, arrows). In contrast, certain sites such as site #1 are bound in all of the assembly states we assayed, suggesting these sites may play critical roles in NC assembly and structure.

### Specific binding of Cp and gRNA site #1 *in vitro*

Cp’s high promiscuity for nucleic acids and anionic molecules *in vitro* induces its assembly into core-like particles (CLPs) that resemble NCs ^20,22,23^. CLP assembly is largely driven by electrostatic interactions with little discrimination among anionic cargoes ^24,39^. Because of this, it is unclear whether specific RNA sequences can drive CLP and NC assembly. To date, no discrete RNA sequence has been shown to be specifically recognized by full length Cp both in infected cells and *in vitro.* Our PAR-CLIP data showed that Cp’s top binding site on the gRNA across all stages of virus assembly was the same sequence (#1 in Fig. 2a, 5c, and 5d). This suggested that site #1 may encompass a distinct RNA motif recognized by Cp. To assess this, we tested whether Cp specifically binds and/or assembles with site #1 *in vitro*. We recombinantly expressed and purified N-terminally Strep-tagged SFV Cp (Fig. S5a). To specifically test binding, Cp was immobilized on Streptactin beads to prevent CLP assembly. Immobilized Cp was then incubated with ^32^P-labeled RNAs representing site #1 or a mutated version identical to that generated in the virus mutant, either alone (first lanes) or in the presence of increasing amounts of an unlabeled random RNA pool (Fig. 6a; upper panels). Both RNAs bound to Cp, but the mutant site #1 RNA was more efficiently competed by random RNA. However, when the binding experiment was performed using Cp preincubated with poly(I:C) to shield non-specific electrostatic interactions, Cp specifically bound site #1 RNA even in the presence of increasing concentrations of random RNA (Fig. 6b; lower panel, left side). In contrast, under the same conditions Cp inefficiently bound the mutant RNA, and its binding was entirely competed by random RNA (Fig. 6b; lower panel, right side). Thus, Cp specifically and directly binds gRNA site #1 both in cells and *in vitro*, demonstrating that site #1 encompasses a motif that Cp specifically recognizes.

**Figure 6.**
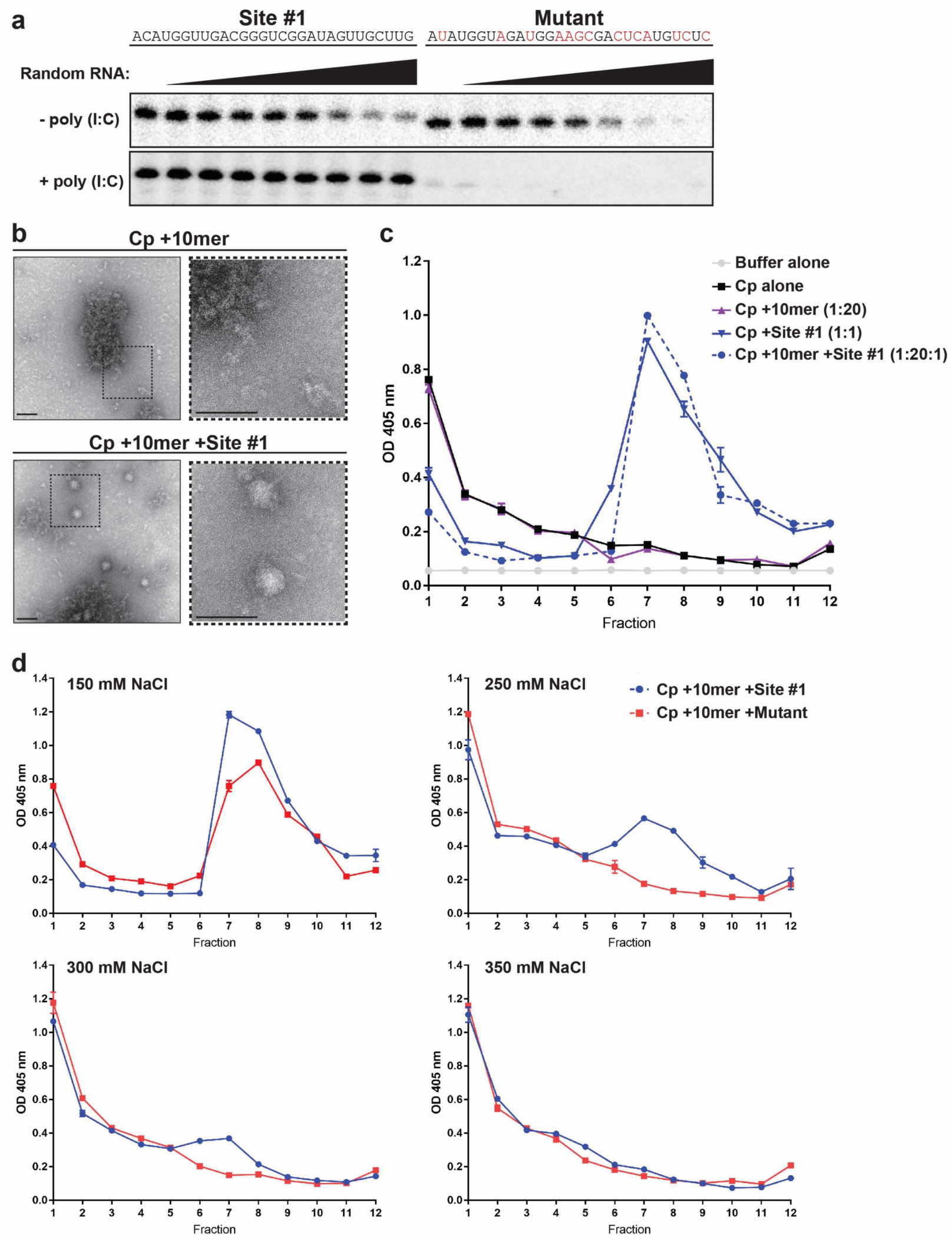
Cp specifically binds and assembles with site #1 *in vitro*. (a) Upper panel: 333 nM ^32^P-labeled site #1 (left) or mutant (right; mutated residues highlighted in red) RNAs were bound to 13.3 pmole Cp immobilized on Streptactin beads in a 30 uL reaction. Increasing concentrations (4-1000 nM) of random RNA were added as a competitor. Bound RNAs were extracted and subjected to Urea-PAGE followed by autoradiography. Lower panel: as above, but immobilized Cp was first preincubated with 20 ug/mL of poly(I:C) to shield non-specific electrostatic interactions. Results are representative of two independent experiments. (b) Negative stain EM of CLP assembly reactions of Cp +10mer or Cp +10mer +Site #1 RNA. Dashed boxes in the right panel are 3X magnification of the corresponding left panel. Scale bar is 80 nm. Images represent one experiment. (c) CLP assembly reactions were incubated for 30 min at 25°C (150 mM NaCl) and then analyzed by sucrose gradient sedimentation. Cp from aliquots of fractions was detected by ELISA. Average absorbances are plotted from duplicate samples with error bars showing the range. Fraction 1 is the top of the gradient. (d) As in (c), except the indicated salt concentrations were used in CLP assembly reactions comparing CLP assembly with site #1 or mutant RNAs. Graphs in (c) and (d) are representative examples of 2-4 independent experiments. See also Figure S5.

We next assessed whether Cp could specifically assemble with site #1 into CLPs. We found that poly(I:C) alone induced CLP formation (data not shown), in keeping with the ability of long anionic polymers such as heparin to promote CLP formation ^24^. Therefore, to shield non-specific electrostatic interactions we instead used a 10 nucleotide DNA oligo (here termed 10mer) that is below the length needed to induce CLP formation ^20^. We confirmed that CLPs did not form with the 10mer by negative stain electron microscopy (EM), which showed amorphous material but no detectable CLPs (Fig. 6b; upper panels). The addition of site #1 RNA produced abundant spherical CLPs of approximately 40 nm in diameter (Fig. 6b; lower), consistent in appearance to that of previously published CLPS ^20,22^. We then quantitated CLP formation by sucrose gradient sedimentation and detection of Cp by ELISA (Fig. 6C and S5b). As predicted, Cp alone or incubated with 10mer (1:20 Cp:10mer) showed most Cp remained unassembled at the top of the gradient (Fig. 6c). Cp incubated with site #1 RNA alone (1:1 Cp:RNA) or in the presence of 10mer DNA (1:20:1 Cp:10mer:RNA) showed a single peak centered toward the middle of the gradient (Fig. 6c), consistent with previous sucrose gradient sedimentation analyses ^20^. Thus, Cp robustly assembles into CLPs with site #1 RNA even in the presence of an excess of electrostatic competitor nucleic acid.

We then tested the effect of mutation of site #1’s sequence on CLP assembly (Fig. 6d). Cp robustly assembled into CLPs with site #1 RNA in the presence of 10mer at physiological salt concentration (Fig. 6d,150 mM NaCl panel). However, under the same conditions CLP formation with the mutant RNA was impaired (Fig. 6d, 150 mM NaCl panel). This decrease in CLP formation was more apparent when the assembly reaction’s salt concentration was raised to shield non-specific electrostatic interactions. Site #1 induced some CLP formation even at 300 mM NaCl, while the mutant only assembled CLPs at 150 mM NaCl (Fig 6d). Thus, site #1 RNA specifically promotes CLP assembly *in vitro* even when general Cp electrostatic interactions are masked, suggesting that site #1 on the gRNA may also act to drive NC assembly in infected cells.

## DISCUSSION

We employed PAR-CLIP and biotin-based Cp retrieval as an experimental approach uniquely suited to define the interactions of the alphavirus Cp with the genomic RNA. Our approach identified a set of high-confidence Cp binding sites on the SFV gRNA, including a novel RNA sequence that specifically bound and assembled with Cp both in infected cells and *in vitro*. The Cp binding sites were preferentially distributed on the first 2/3 of the gRNA, suggesting a molecular rationale for the previously observed selective packaging of gRNA over sgRNA ^9–11^. An overall preferential binding to gRNA-specific sequences was maintained across different NC assembly states. Such continuous engagement with gRNA-specific sequences can promote gRNA packaging and high infectivity.

A previous study used CLIP to identify SINV Cp interactions with the gRNA in infected cells ^12^. Comparison with our results showed that the SINV and SFV Cp top binding sites do not overlap, and that SINV Cp’s major binding sites are located within the last ∼1/3^rd^ of the gRNA sequence, overlapping with the sgRNA. Mutation of the SINV Cp binding sites does not affect gRNA packaging or NC assembly, but instead affects the stability of the incoming gRNA. These differences may reflect biological differences between SINV and SFV, but could also be due to differences in experimental approaches.

Alphavirus gRNA packaging signals were originally identified through analysis of DI-RNAs ^18,19,40^. Although elegant studies showed that appending the SINV PS onto reporter or helper RNAs could increase their packaging in replicon systems and their direct binding to Cp *in vitro*, the increase was at most 5-fold ^18,41^. Conversely, SINV or VEEV with mutated PSs are impaired but viable ^17^. While our data are the first to show that the alphavirus Cp directly binds the DI-RNA-derived PS in cell culture, we also found that genome packaging within the SFV complex does not require the PS. Instead, our data argue that packaging involves preferential Cp binding to multiple gRNA-specific sites. Although the relative importance of the PS appears to vary between alphavirus complexes, it seems clear that the PS is not strictly required for gRNA packaging/NC assembly. Given that recent studies indicate that DI-RNAs and defective viral genomes can have multifaceted functions (reviewed in ^42^), the prior DI-RNA studies may have more complex interpretations than previously suspected.

We found that while mutation of multiple high affinity Cp binding sites did not significantly affect viral replication or protein expression, it significantly reduced infectious virus production. This correlated with reduced gRNA but elevated sgRNA levels in virus particles. Inhibition of infectious virus production was strongest at early stages of virus assembly, in keeping with increasing Cp concentrations at later times of infection driving Cp engagement with mutated binding sites. Based on our results we propose a mechanism for selective genome packaging via Cp’s interactions with multiple sites selectively located on the gRNA (Fig. 7). At early stages of virus assembly, gRNA levels are high but intracellular Cp levels are relatively low, leading to Cp binding only to high affinity sites on the gRNA (Fig. 7, ia vs. ib). This stage may involve Cp dimers ^21^, and could be represented by a proposed 90S NC assembly intermediate, which has a lower Cp:gRNA ratio than cellular NC ^43–45^. For some alphaviruses such as SINV and VEEV, this early stage may involve strong engagement of Cp with the PS. As Cp levels accumulate in the cell, Cp begins engaging with lower affinity sites on the gRNA, enabling compaction of the gRNA molecule through charge neutralization ^15,37,46^ (Fig. 7, ii). More Cp would then be recruited to the assembling NC ^47^ (Fig. 7, iii), and Cp-Cp interactions and E2-Cp interactions would impose T=4 icosahedral symmetry and promote budding, resulting in formation of the completed viral NC and Cp-RNA interactions (Fig. 7, iv). Additional unknown factors may influence genome packaging and NC assembly, including specific subcellular locations or host factors.

**Figure 7.**
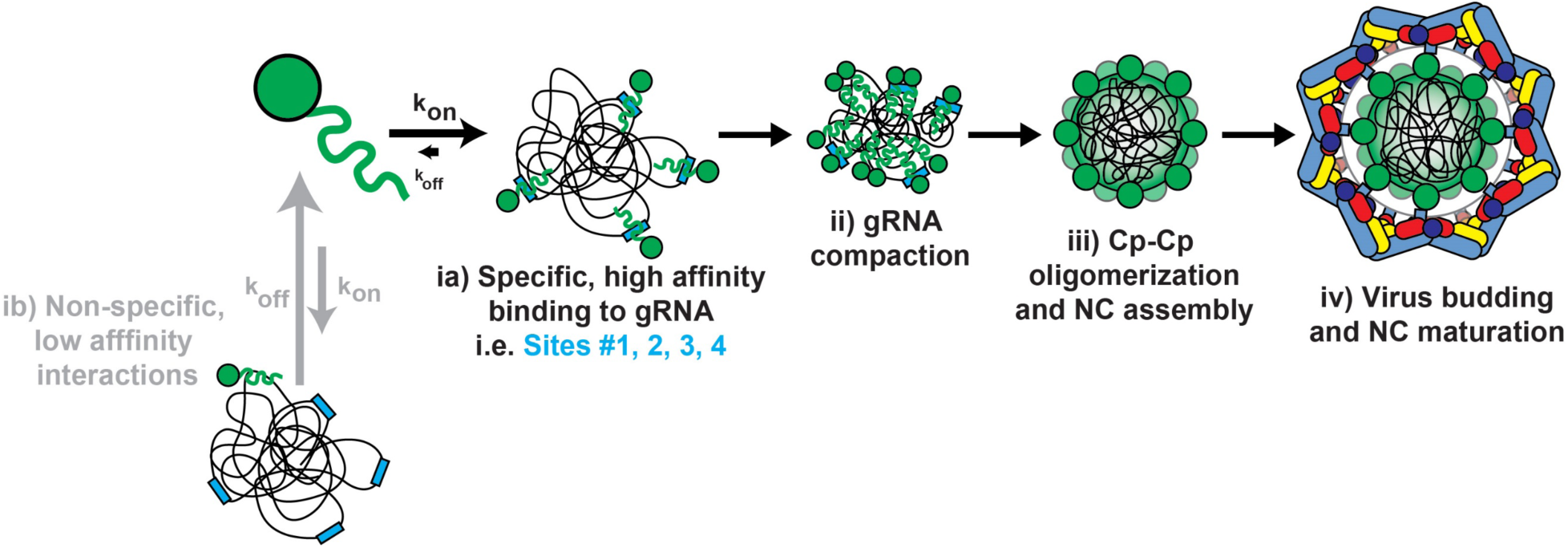
A multi-site genome packaging model for alphaviruses. ia) Cp (green) specifically interacts with its high affinity sites (blue rectangles) on the gRNA (black lines) and stably binds them. ib) Cp can also non-specifically interact with low-affinity sites on the gRNA or other RNAs such as the sgRNA or cellular RNAs (not shown), but binding is not stable. ii) As Cp levels increase in the cell, Cp continues to engage multiple high affinity sites on the gRNA, enabling genome compaction. iii) Further Cp recruitment leads to Cp oligomerization and full NC assembly. iv) Virus assembly and budding cause discrete changes in Cp:gRNA contacts within the viral NC.

From a broader perspective, a multi-site packaging mechanism could be favorable for many RNA viruses given that RNA polymerases generally have low fidelity. Their relatively high mutational frequency produces genetic diversity important for RNA virus adaption to environmental changes, but can also decrease virus fitness ^48,49^. A multi-site packaging mechanism could help to ensure correct genome packaging even if deleterious mutations arose in several Cp binding sites. For example, retroviruses such as HIV-1 have high mutation rates and a defined packaging signal known as the Ψ element (reviewed in ^50^). However, Ψ mutants only modestly decrease (∼3-5 fold) infectious virus production or genome packaging ^32,51^. Multi-site packaging mechanisms have been described for RNA viruses such as parechovirus 1 ^52^ and hepatitis B virus ^53^, as reviewed in ^54^. In addition, multi-site packaging may promote successful co-assembly of Cps and large ssRNAs while decreasing non-productive assembly pathways ^38^. Thus, the multi-site-based packaging that we describe here for alphaviruses may be representative of a more general mechanism utilized by many RNA viruses to selectively package their genomes.

Our data represent the first demonstration of a discrete RNA sequence, site #1, with which alphavirus Cp specifically binds and assembles both in cells and *in vitro*. While multiple binding sites such as site #1 were maintained across virus assembly, we also observed discrete differences in Cp:gRNA interactions between the cellular and viral NCs. This could reflect changes induced by different chemical environments, or could be induced by envelope protein association and virus budding. Changes in NC architecture during various stages of virus exit have been previously observed using morphological and biochemical techniques ^55–58^. Our data thus suggest that such changes in the NC structure also significantly affect Cp-gRNA binding. If E2 interactions/budding affect Cp-gRNA interactions, this would suggest that changes in NC architecture are propagated from the outer, envelope-protein interacting face of the NC shell to the inner RNA core of the virus particle. Further functional analyses are needed to determine the potential importance of such changes in virus Cp-RNA interactions during exit.

## Author contributions

R.S.B. and M.K. conceived the project. R.S.B. and D.G.A. made the PAR-CLIP libraries. D.G.A. performed the *in vitro* binding assay. R.S.B. performed all other experiments. M.K. and M.H. supervised the research. R.S.B. and M.K. wrote the original draft. R.S.B., D.G.A., M.H., and M.K. reviewed and edited the draft.

## METHODS

### Cells

Baby hamster kidney (BHK-21) cells (gift from Dr. Ari Helenius) were cultured in Dulbecco’s modified Eagle’s medium (DMEM, HyClone) containing 4 mM L-glutamine, 100 U penicillin/mL, 100 μg streptomycin/mL, 10% tryptose phosphate broth, and 5% FBS at 37°C. Vero cells (ATCC; gift from Dr. Kartik Chandran) were cultured in DMEM containing 4 mM L-glutamine, 100 U penicillin/mL, 100 μg streptomycin/mL, and 10% FBS at 37°C. Vero cells stably expressing HA-BirA were generated by transfecting cells with the expression plasmid described below using Lipofectamine 2000 (Invitrogen). 48 h post-transfection, cells were split to 25% confluency and selected with 1 mg G418/mL (Sigma-Aldrich) for two weeks. G418-resistant cells were then seeded sparsely and individual clones isolated using cloning cylinders (Corning). Clones were screened for their ability to biotinylate Cp-mAVI by western blot and fluorescence microscopy using streptavidin probes. Cell lines were authenticated by morphologic evaluation and were checked for mycoplasma contamination (MycoAlert^TM^ PLUS Mycoplasma Detection Kit, Lonza).

### Antibodies

The following antibodies and SA probes were used for western blots and immunofluorescence as indicated: Rabbit polyclonal anti-α tubulin (Abcam, 18251), Rabbit polyclonal anti-E2/E1 ^60^, monoclonal anti-Cp (42-1) ^61^, monoclonal anti-E2 (E2-1) ^62^, Streptavidin Alexa Fluor 680 Conjugate (ThermoFisher Scientific, S32358). Rabbit polyclonal anti-nsP2 was a gift from Dr. Andres Merits. Rabbit polyclonal anti-Cp was generated (Covance) against the C-terminal domain of SFV Cp (residues 119-267) expressed and purified as described below for full-length Cp. Specificity was confirmed by western blot, immunofluorescence of uninfected vs. infected cells, and co-staining with monoclonal antibody 42-1.

### Viruses, mutants, and plasmids

A minimal biotin acceptor motif (mAVI tag; LNDIFEAQKIEWH) ^29,30^ was inserted after Cp residue 95 by overlapping PCR using restriction sites XbaI and NsiI in the SFV infectious clone pSP6-SFV4 ^63^. Overlapping PCR primers: 5’CTGAACGACATCTTCGAGGCCCAGAAAATCGAATGGCACaagcaagccgacaag3’ and 5’GTGCCATTCGATTTTCTGGGCCTCGAAGATGTCGTTCAGgtctttcttcttttgctgc3’. Individual Cp binding site mutants were constructed by overlapping PCR and using the nearest unique restriction sites present in pSP6-SFV4. Double and Quintuple Cp binding site mutants were constructed by restriction digest and subcloning from individual mutants. The 17-site mutant, Full PS mutant, and 17-site + Full PS mutant were custom synthesized (Epoch Life Science, Inc), and subcloned by restriction digest into pSP6-SFV4. The Chikungunya Full PS mutant was custom synthesized (Epoch Life Science, Inc) and subcloned by restriction digest into the pSinRep5-181/25 Chikungunya infectious clone (kindly provided by Dr. Terrence Dermody ^64^). All infectious clone mutants were verified by sequencing the entire region affected by the cloning approach (Genewiz, South Plainfield, NJ) and analyzed by restriction digest to check for plasmid rearrangements. A humanized BirA construct was provided by Vasily Ogryzko ^65^ and subcloned into pcDNA3.1+ with an N-terminal HA-tag added by PCR.

### Virus stocks

SFV WT and mAVI virus stocks were generated by electroporating BHK cells with *in vitro* transcribed (IVT) viral RNA and collecting the cell media at 24 h post- electroporation ^63^. Virus-containing media were clarified by centrifugation at 10,621 X g 4°C for 10 min, and 10 mM HEPES pH 8.0 was added to the supernatant before aliquoting and freezing. Virus stocks for growth comparisons of SFV WT, Full PS mutant, and the indicated Cp binding site mutants were generated the same way except that the cell media were collected at 8 h post-electroporation. CHIKV WT and Full PS mutant stocks were generated as above except that the cell media were harvested at 22 h post-electroporation. All virus stocks were titered in two independent experiments by plaque assay on BHK cells.

### Virus growth curves

Growth curves were performed on Vero cells infected at the indicated multiplicity of infection (MOI) for 1.5-2 h at 37°C. At the indicated time points, the virus-containing media were collected, clarified, aliquoted and frozen at -80°C. Aliquots were titered via plaque assay on BHK cells.

### Cell lysis and western blot

Vero parental or Vero+BirA cell lines were infected at an MOI=10 for 1.5 h at 37°C before transfer into fresh medium containing 50 µM biotin. At the indicated time points, the cells were washed and lysed with lysis buffer [50 mM Tris-Cl pH 7.4, 100 mM NaCl, 1% Triton-X-100, 1 mM EDTA, 6 mM NaPPi (to inhibit post-lysis biotinylation), and an EDTA-free protease inhibitor cocktail (Roche; 1 tablet/10 mL)] on ice. The lysate was then clarified by centrifugation and the soluble lysate was frozen at -80°C. Lysates were subjected to SDS-PAGE followed by transfer to nitrocellulose membranes. Membranes were probed with the indicated primary antibodies and corresponding secondary antibodies conjugated to Alexa Fluor 680 or 800 dyes before imaging on an Odyssey Fc Imaging System (LI-COR Biosciences).

### Immunofluorescence

Vero parental or Vero+BirA cells were infected at an MOI=1 for 1.5 h at 37°C, and then fresh medium supplemented with 50 µM biotin was added to each well. At 7 hpi, the cells were fixed with 4% paraformaldehyde (Electron Microscopy Sciences) and quenched with 50 mM NH_4_Cl. The cells were permeabilized with 0.1% Triton-X-100 and blocked with 0.2% gelatin. Coverslips were then stained with the indicated primary antibodies followed by the corresponding secondary antibody conjugated to an Alexa- Fluor dye. Images were acquired on a Zeiss Axiovert 200 M microscope and processed using ImageJ.

### Transmission electron microscopy

Vero parental and Vero+BirA cell lines were infected with SFV WT or mAVI at an MOI=10 for 1.5 h at 37°C, before supplementing with fresh medium containing 50 µM biotin. At 7.5 hpi, the cells were washed once with serum-free medium and then fixed with 2.5% glutaraldehyde and 2% paraformaldehyde in 0.1 M sodium cacodylate buffer for 30 minutes at room temperature. The Einstein Analytical Imaging Facility then processed the samples by postfixing with 1% osmium tetroxide and 2% uranyl acetate. The samples were dehydrated and embedded, and thin sections were stained with uranyl acetate followed by lead citrate. Images were taken on a JEOL 1200EX electron microscope at 80 kV and assembled using Adobe Photoshop software. For negative stain analysis of CLP assembly reactions, 5 µL of sample was applied to glow-discharged carbon-coated copper grids and stained with 1% v/v uranyl acetate. Grids were air dried and then viewed on a FEI Tecnai 20 electron microscope at 120 kV.

### Specific Infectivity

Vero parental and Vero+BirA cells were infected at an MOI=10 for 1.5 h at 37°C before supplementing with fresh medium containing 50 µM biotin. At 7.5 hpi, the virus-containing media were collected and clarified by centrifugation. An aliquot was taken from the clarified supernatant for titering by plaque assay to determine infectious particle number. The remaining supernatant was layered over a 20% sucrose cushion and centrifuged at 35,000 rpm (Beckman Coulter SW41 rotor) for 3 h at 4°C. The pelleted virus was resuspended in TN buffer [50 mM Tris-Cl pH 7.4, 100 mM NaCl, and a protease inhibitor cocktail (Roche; 1 tablet/10 mL)] on ice overnight. Virus suspensions were serially diluted, boiled in SDS sample buffer, and subjected to SDS-PAGE followed by western blot using an E2-specific monoclonal antibody. E2 signal was imaged and quantified using the Odyssey Fc Imaging System (LI-COR Biosciences) to determine particle number. The specific infectivity was calculated as a ratio of the infectious particle number (plaque forming units) to the E2 signal (total particle number).

### PAR-CLIP

PAR-CLIP on SFV Cp was performed based on previously described methods ^66–69^.

#### Total cellular Cp population

Vero+BirA cells were counted and seeded in 35 mm dishes 24 h before infection. The cells were infected with SFV mAVI at an MOI=10 for 1.5 h, washed three times, and then placed into media supplemented with 50 µM biotin and 100 µM 4-thiouridine. At 7 hpi, the cells were UV-irradiated with 368 nm light at 0.15 J/cm^2^, or control cells were mock UV-irradiated. Cells were then placed on ice, washed three times with PBS, and then lysed with lysis buffer (50 mM Tris-HCL pH 7.4, 100 mM NaCl, 1% NP-40, 1 mM EDTA, 6 mM NaPPi, 0.5 mM DTT, and a complete protease inhibitor tablet (Roche; 1 tablet/10 mL)) for 10 min with gentle rocking. The lysate was clarified by centrifugation at 13,000 rpm 4°C for 10 min. The soluble lysate was then treated with RNaseT1 (1 U/uL) for 15 min at room temperature and then placed on ice for 5 min. The samples were then frozen at -80°C and a 5% input control was taken at this step to compare Cp expression levels by western blot (Fig. S2c). Streptavidin Dynabeads (SA-DB; Dynabeads MyOne Streptavidin C1, ThermoFisher Scientific) were equilibrated with PBS while samples thawed on ice. A final concentration of 0.5% SDS was added to the samples, mixed, and then incubated with the SA-DB by rocking at 4°C for 3 h. The SA-DB were then washed twice with RIPA buffer supplemented with high salt (10 mM Tris-Cl pH 7.4, 0.5 M NaCl, 1 mM EDTA, 1% NP-40, 1% sodium deoxycholate, 0.1% SDS, 0.5 mM DTT, and a complete protease inhibitor tablet (Roche; 1 tablet/10 mL)) for 5 minutes per wash, once with PBS, and treated with RNaseT1 at 1 U/μL for 15 min at room temperature before cooling on ice for 5 min. The SA-DB were washed again with high salt RIPA buffer for 5 min, then washed twice with dephosphorylation buffer (50 mM Tris-Cl pH 7.9, 100 mM NaCl, 10 mM MgCl_2_, and 1 mM DTT), and treated with calf-intestine phosphatase at 0.5 U/μL for 10 min at 37°C. The SA-DB were washed once with lysis buffer, twice with PNK buffer (50 mM Tris-Cl pH 7.5, 50 mM NaCl, 10 mM MgCl_2_, 5 mM DTT) lacking DTT, then resuspended in complete PNK buffer. 1 U/μL of T4 PNK and 0.5 μCi/uL of ^32^P-ATP were added to the SA-DB for 30 min at 37°C with gentle mixing every 5 min. After 30 min, 100 µM cold ATP was added for 5 min at 37°C. The SA-DB were washed five times with PNK buffer without DTT and then once with PBS. Samples were boiled for 5 min in SDS sample buffer four sequential times to maximize elution before pooling eluates. The eluates were subjected to SDS-PAGE and then transferred to nitrocellulose membranes for autoradiography. The film was overlaid on top of the membrane and the corresponding CpxRNA crosslinked adducts were excised from the membrane. The membrane was treated with proteinase K for 90 minutes at 55°C, and the RNA was extracted with phenol-chloroform and precipitated.

#### Cellular and viral NCs

PAR-CLIP on cellular and viral NCs was performed as described above, but with the following differences. Two 10 cm plates of Vero+BirA cells were infected with SFV mAVI at an MOI=10. At 7.5 hpi, the virus-containing media were collected and clarified by centrifugation at 10,000 rpm for 5 min at 4°C. The virus was then purified over a 20% sucrose cushion by centrifugation at 35,000 rpm (Beckman Coulter SW41 rotor) for 3 h at 4°C. The pellet was resuspended on ice for 3 h in TN buffer and then transferred to a 35 mm dish. The virus was UV irradiated or mock irradiated with 368 nm light at 0.15 J/cm^2^, and then lysed in lysis buffer, denatured with 0.5% SDS, and processed as described under *Total cellular Cp population*. The cells from the same plates were washed with PBS and then UV-irradiated with 368 nm light at 0.15 J/cm^2^ or mock-irradiated. Cells were pelleted, lysed in lysis buffer, clarified by centrifugation at 13,000 rpm for 10 min at 4°C, and then treated with 25 mM EDTA for 20 min to dissociate polysomes. Samples were loaded onto 7.5-20% (wt/wt) sucrose gradients in TN buffer + 2 mM EDTA and 0.1% NP-40, and centrifuged at 41,000 rpm (Beckman Coulter SW41 rotor) for 2 h at 4°C. 1 mL fractions were collected and aliquots were analyzed by SDS-PAGE followed by western blot using an anti-Cp antibody to identify the NC fractions. The NC fractions were pooled, denatured with 0.5% SDS, retrieved with SA-DB, and processed as described under *Total cellular Cp population*.

### cDNA library construction and sequencing

The extracted RNA was ligated to the 3’ adenylated adapter using Rnl2(1-249)K227Q ligase at 4°C overnight. A size marker mix containing synthetic 19 and 35 nt long RNAs was used as a positive control for all ligation steps. After ligation, the reaction was denatured at 90°C for 1 min and electrophoresed on a 15% denaturing Urea-PAGE. The gel was then exposed to a phosphoimager screen for an hour at -20°C. The image was printed to its original size and aligned on to the gel where the successfully ligated product was excised, shredded, and incubated with 0.3 M NaCl at 60°C for 45 minutes. The sample was filtered and then the RNA was precipitated in ethanol at -20°C for at least an hour. The precipitated RNA was pelleted and dissolved in water. The 5’ adapter was ligated to the sample using Rnl1 for 1 h at 37°C. The sample was then processed the same way as the 3’ adapter ligation reaction. The ligated sample was reverse transcribed to make cDNA using Superscript III reverse transcriptase for 2 h at 50°C. A pilot PCR was performed to optimize PCR cycle number to prevent over amplification of the library. This entailed taking a 10 uL aliquot from the pilot PCR after every three cycles between cycles 12 and 30 for agarose gel analysis. The lowest PCR cycle number to generate a visible PCR product by ethidium-bromide staining was used for the final PCR. The final PCR product was purified away from linker-linker (3’ adapters ligated to 5’ adapters) byproducts using a 3% Pippin Prep (Sage Science) before sequencing on an Illumina MiSeq machine. Oligonucleotides used (5’ to 3’):

RNA PCR Index Primer 9

CAAGCAGAAGACGGCATACGAGATCTGATCGTGACTGGAGTTCCTTGGCACCCGA

GAATTCCA;

RNA PCR Index Primer 10

CAAGCAGAAGACGGCATACGAGATAAGCTAGTGACTGGAGTTCCTTGGCACCCGA

GAATTCCA;

RT Primer GCCTTGGCACCCGAGAATTCCA;

3’ Barcode adapter 29.31 5’-rAppNNTAGCGATGGAATTCTCGGGTGCCAAGG-L;

3’ Barcode adapter 29.32 5’-rAppNNCTGTAGTGGAATTCTCGGGTGCCAAGG-L;

3’ Barcode adapter 29.33 5’-rAppNNTAGTCGTGGAATTCTCGGGTGCCAAGG-L;

L = aminolinker, 3’-amino modifier C7.

### RT-qPCR

To quantify viral RNA in infected cells, Vero cells were mock infected or infected with SFV WT or 17-site mutant at an MOI=0.01 for 1.5 h at 37°C. At the indicated time points, cell-associated RNA was extracted using Trizol as per the manufacturer’s instructions. Total RNA was quantified by Nanodrop and RNA integrity was assessed by 2% (v/v) bleach agarose gel electrophoresis ^70^ to visualize rRNA. To quantify viral RNA in released particles, Vero cells in two 10 cm plates were infected as above. At the indicated time points, virus-containing media were collected and clarified by centrifugation. An aliquot was taken from the clarified supernatant for titering via plaque assay to determine infectious particle number. The remaining supernatant was layered over a 20% sucrose cushion and pelleted as previously described (c.f. Methods, Specific Infectivity). Pellets were resuspended in 100 µL of 50 mM Tris pH 7.4, 100 mM NaCl, 1 mM EDTA, protease inhibitor (Roche; 1 tablet/10 mL), 1 U/uL recombinant Rnasin (RNase inhibitor) before extracting viral RNA using the MagMax viral RNA isolation kit as per the manufacturer’s instructions. IVT gRNA was produced from SFV infectious clone plasmid and treated with 10 U of DNase1 (RNase-Free; NEB) for 30 min at 37°C before purification with the RNeasy MiniElute Cleanup kit (Qiagen) as per the manufacturer’s instructions. IVT gRNA integrity and purity was assessed by agarose gel analysis and quantified by NanoDrop. cDNA was synthesized from 5 ng of extracted cell-associated RNA, 5 ng of IVT gRNA, or 10% of pelleted viral RNA using the Verso cDNA synthesis kit and gRNA-specific or [gRNA+sgRNA] reverse primers. qPCR was performed using Power SYBR Green PCR Master Mix (Applied Biosystems) in a ViiA 7 Real-Time PCR machine using 384-well plates. cDNA from samples corresponding to three biological replicates were assayed with technical duplicates. A no template control and mock-infected controls were used to ensure viral RNA-specific amplification. Four standard curves consisting of a 10-fold dilution series of IVT gRNA cDNA were used to calculate sample gRNA and [gRNA+sgRNA] copy numbers by relating the threshold cycle values. Cycle conditions were 50°C 2 min, 95°C 10 min, and 45 cycles of 95°C 15 s and 60°C 1 min, followed by melt curve analysis. Oligonucleotides used (5’ to 3’): gRNA-specific forward primer CTACGCTACACCAGATGAATACC, gRNA-specific reverse primer GGCTATGTCTGCTCTCTTAACTC, [gRNA+sgRNA] forward primer GCCTCGAACCAACCCTTAAT, and [gRNA+sgRNA] reverse primer CTTCTCTTTAGTGGAGCACTCTG.

### Recombinant Cp expression and purification

SFV Cp was cloned via PCR into pET29a with an N-terminal double Strep Tag (WSHPQFEK) followed by a glycine+serine linker and a TEV protease cleavage site (2XStrep_GS_TevClvg_Cp) for expression in Rosetta2 cells. Rossetta2 cells were grown at 30°C until an OD_600_ of 1.0, and protein expression was induced with IPTG at 16°C overnight. Cells were harvested and resuspended in binding buffer (100 mM Tris- Cl pH 8.0, 150 mM NaCl, 1 mM EDTA, complete protease inhibitor cocktail (Roche; 1 tablet/10 mL) before sonicating on ice. The lysate was clarified by two centrifugation steps. The soluble lysate was then diluted to 10X the volume in binding buffer supplemented with 1.5 M NaCl, mixed, and placed on ice overnight. The next day, the protein was purified via affinity chromatography using Strep-Tactin sepharose (IBA) and then dialyzed into 50 mM Tris-Cl pH 7.4 and 100 mM NaCl before concentrating and freezing. Protein purity (>99%) was assessed by SDS-PAGE and Coomassie staining (Fig. S5a). The sample’s A_260_/A_280_ ratio showed no RNA or DNA contamination.

### *In vitro* core-like particle (CLP) assembly

HPLC purified RNA oligos corresponding to site #1 or its mutant version were purchased from IDT. RNAs were denatured at 90°C for 2 min and chilled on ice for 10 min before serially diluting to 10 µM in 50 mM K-HEPES pH 7.0, 200 mM KCl, 10 mM MgCl_2_. Diluted RNAs were incubated at 37°C for 30 min to promote folding before placing back on ice. CLPs were assembled in a 100 uL volume in reaction buffer [50 mM Tris pH 7.0, 150 mM NaCl, 5 mM MgCl_2_, 5 mM KCl, 0.01% (v/v) Tween 20, 1 µg/µL BSA, 5 mM DTT, and 0.8 U/µL of recombinant RNasin (RNase inhibitor)]. Cp was diluted (500 nM final concentration) in reaction buffer before adding 10mer DNA oligo (CCGTTAATGC; 10 µM final concentration) or buffer control (50 mM Tris pH 7.4, 100 mM NaCl) for 10 min at RT. Reactions were placed on ice and then each RNA oligo (500 nM final concentration) or buffer control was added. CLP assembly reactions were incubated at 25°C for 30 min before loading onto 15-30% (wt/wt) sucrose gradients and spinning at 159,599 X g for 43 min at 4°C (Beckman Coulter TLS 55 rotor). Gradients were immediately fractionated and aliquots of each fraction were denatured by boiling in 1% SDS to ensure antibody access, and then diluted into binding buffer (50 mM Tris pH 8.0, 150 mM NaCl, 2 mM EDTA, and 1% NP-40) to make a mixed detergent micelle (∼0.075% SDS final concentration). A corresponding standard curve consisting of a 2- fold dilution series of purified Cp was similarly processed to mimic sucrose, SDS, and boiling conditions (Fig. S5b) for every CLP experiment. Samples were bound in duplicate or triplicate to high-capacity streptavidin-coated plates (Pierce) preblocked with 1% BSA in TBS (25 mM Tris pH 7.2, 150 mM NaCl) for ∼12-18 h at 4°C. Plates were incubated with a polyclonal antibody to Cp followed by secondary antibody conjugated to alkaline phosphatase. pNPP substrate was added for 15 min before measuring absorbance at 405 nm on a Tecan Infinite F50 plate reader.

### *In vitro* binding assay

*In vitro* binding assays were performed following the RNA Bind-n-Seq protocol ^71^ with the following modifications. Recombinant Cp was immobilized on Strep-TactinXT coated magnetic beads (20 pmole protein/μl bead slurry) in buffer w (100 mM Tris-Cl pH 8.0, 150 mM NaCl, 1 mM EDTA). Beads were resuspended in RNA binding buffer (25 mM Tris pH 7.5, 150 mM KCl, 3 mM MgCl_2_, 0.01% (v/v) Tween 20, 1 mg/mL BSA, 1 mM DTT). Binding reactions were prepared in 96 well plates. A total of 13.3 pmole of Cp was immobilized on beads per 30 μl reaction and concentrations of 333 nM site #1 or its mutant along with increasing concentrations of random 29 mer RNA (4-1000 nM) were added in each well with or without 20 μg/ml poly(I:C) preincubation. Binding was performed for 30 minutes at 30°C and the plate was then placed on a 96 well magnetic rack. The beads were washed 3 times with wash buffer (25 mM Tris pH 7.5, 150 mM KCl, 60 ug/mL BSA, 0.5 mM EDTA, 0.01% (v/v) Tween 20). 40 μl of elution buffer (10mM Tris pH 7.0, 1mM EDTA, 1% SDS) was added to the beads before heating at 70° C for 10 minutes. The supernatant from each well was placed in a new plate and the bound RNA was purified away from Cp by phenol/chloroform isolation. The purified RNA was resolved by 15% denaturing Urea-PAGE.

### PAR-CLIP data analysis

Adapters were removed using Cutadapt ^72^ and the remaining sequences were mapped to the SFV4 genome using Bowtie. Only mapped sequences that contained the characteristic T-to-C mutation were further used. Initial analyses compared sequence reads (uncollapsed reads) vs. unique reads stemming from the barcoded adapters (collapsed reads) to assess PCR duplicates. No significant difference was observed in Cp’s top binding sites between the uncollapsed vs. collapsed reads, which was expected because we carefully optimized PCR cycle number during cDNA library construction (as described above). We therefore continued our analyses with the uncollapsed reads. Reads were normalized to the nucleotide position with the highest read frequency within the library, and then biological replicates were averaged and renormalized to produce the final normalized read density. High confidence binding sites were manually defined with the following criteria: the binding site must be at least 10 nt long, have at least one position containing an average normalized read density ≥5.0, and both replicate libraries must have ≥ 250 reads at that position. Within these binding sites, the 5’ and 3’ ends were generally defined by having ≥1.5 average normalized read density, ≥10 reads per replicate library, and 3’ G-bias due to RNaseT1 digest. Low confidence binding sites were defined as those with at least an average normalized read density ≥1.0 and ≥10 reads for each replicate library.

### Statistical analysis

Statistical analyses were performed using GraphPad Prism 7.04 (GraphPad Software). The specific statistical tests used and the number of replicates per experiment are stated in the figure legends. In all graphs, four asterisks indicate a P-value of <0.0001, three asterisks indicate <0.001, two asterisks indicate <0.01, one asterisk indicates <0.05, and ns indicates >0.05.

## DATA AVAILABILITY

Upon publication, the datasets generated during and/or analyzed during the current study will be made available in the NCBI GEO repository.

## Acknowledgements

We thank all of the members of the Kielian lab and Dimitrios Zattas for helpful discussions and comments on the manuscript. We thank Matthew Angeliadis for technical assistance. We thank the Einstein Analytical Imaging Facility and facility members Xheni Nishku and Timothy Mendez for training and technical assistance on the electron microscope. We thank Susan Buhl and Matthew Scharff for technical assistance and use of their plate reader.

This work was supported by grants to M.K. from the National Institute of General Medical Sciences (R01-GM057454) and the National Institute of Allergy and Infectious Diseases (R01-AI075647) and by Cancer Center Core Support Grant NIH/NCI P30- CA13330. R.S.B. was supported by an NIH NRSA (F32-GM122450) postdoctoral fellowship and the Charles H. Revson Senior Fellowship in Biomedical Sciences. M.H. and D.G.A are supported by the Intramural Research Program of the National Institute of Arthritis and Musculoskeletal and Skin Diseases. The content of this paper is solely the responsibility of the authors and does not necessarily represent the official views of the National Institute of General Medical Sciences, the National Institute of Allergy and Infectious Diseases, National Institute of Arthritis and Musculoskeletal and Skin Diseases, or the National Institutes of Health.

